# Selfing Maintains Flower Colour Polymorphism in *L. arvensis* Despite High Inbreeding Depression

**DOI:** 10.1101/761122

**Authors:** Francisco Javier Jiménez-López, Pedro Luis Ortiz, María Talavera, Montserrat Arista

## Abstract

Flower colour polymorphism (FCP) is frequently associated with differences in pollinator attraction. FCP maintenance is intriguing, as positive directional selection by pollinators should result in the loss of polymorphism. Autonomous selfing could confer reproductive assurance when pollen is limited, and could be a mechanism for maintaining polymorphism unless inbreeding depression is high. We study the role of selfing in maintaining FCP in *Lysimachia arvensis*, a species with blue and red morphs co-occurring in Mediterranean populations, where pollinators negatively select for the red morph. We experimentally assessed inbreeding depression in both morphs in two Mediterranean populations and genetic diversity was studied via AFLP and SSR microsatellites in 20 populations. Between-morph genetic differentiation was high and the red morph had a lower genetic diversity, mainly in the Mediterranean. Results also show strong phenological differences between selfed and outcrossed progeny, and a high ID of the red morph. The low genetic diversity of the red morph is in concordance with a reproductive system based predominantly on selfing. However, ID suggests a limited capacity for red morph recruitment, according to its low frequency in polymorphic populations. Genetic differentiation between morphs indicates a low gene flow between them, opening the possibility of reproductive isolation and speciation in *Lysimachia arvensis*.

## Introduction

Flower colour polymorphism (FCP) is defined as “*the presence of at least two genetically-determined colour morphs within a single interbreeding population, the rarest of which is too frequent to be solely the result of recurrent mutation”* (Huxley, 1955). Flower colour polymorphism is a widespread trait in plants, but is relatively infrequent (e.g. Whitney, 2005; Rausher, 2008; Narbona, Wang, Ortiz, Arista, & Imbert, 2018). In many cases, both biotic and abiotic factors influence the fitness of plants with flowers of a certain colour. Thus, floral morphs can show differential tolerance to abiotic factors (Warren & Mackenzie, 2001; Arista, Talavera, Berjano, & Ortiz, 2013) due to the association between some floral pigments and protective flavonoids. Similarly, biotic agents of selection such as herbivores (Strauss & Whittall, 2006; Sobral et al., 2015) or pollinators (e.g. Meléndez-Ackerman, Campbell, & Waser, 1997; Jones & Reithel, 2001) can show a preference for a particular morph, thereby being responsible for its lower or higher fitness, respectively. Under these circumstances, the maintenance of flower colour polymorphism is an apparent enigma, as directional selection along with genetic drift should drive the loss of one of the colour morphs.

Pollinators are among the most important biotic factors involved in flower colour selection (e.g. Fenster, Armbruster, Wilson, Dudash, & Thomson, 2004; Whibley et al., 2006; Wessinger & Rausher, 2012; Schiestl & Johnson, 2013). They usually discriminate between colour morphs and can show preferences for a particular colour causing positive (e.g. Morgan & Schoen, 1997; Jones & Reithel, 2001; Ortiz, Berjano, Talavera, Rodríguez-Zayas, & Arista, 2015), or negative directional selection (e.g. Waser & Price, 1981, 1983; Gigord, Macnair, & Smithson, 2001; Kagawa & Takimoto, 2016). In polymorphic species, the less-visited morph could suffer a fitness reduction leading to negative directional selection, which along with genetic drift should result in the loss of polymorphism and the evolution of monomorphic populations (Waser & Price, 1981; Levin & Brack, 1995; Campbell, Waser, & Meléndez-Ackerman, 1997; Jones & Reithel, 2001). However, the less-visited morph will suffer fitness reduction only if it depends strictly on pollinators to reproduce. Some plants have the capacity to produce seeds by autonomous selfing when pollinators are scarce, thereby showing a mixed mating system. This capacity, called reproductive assurance (RA), allows plant reproduction when opportunities for outcrossing are reduced (Holsinger, 2000), as occurs when pollinator attention is low.

Plants with reproductive assurance capacity via selfing can be independent of pollinators and can be maintained in populations, at least for some time (Takebayashi & Morrell, 2001; Charlesworth, 2006). However, the long-term genetic consequences of selfing are largely known. Mating system, which is responsible for gene transmission between plants and generations, is one of the most important factors affecting the pattern of gene diversity of plant populations (Loveless & Hamrick, 1984; Hamrick & Godt, 1996; Glémin, Bazin, & Charlesworth, 2006). Outcrossing is more advantageous than selfing, because it maintains higher levels of gene diversity in populations, which should increase their adaptive potential. In fact, selfing causes a reduction of gene diversity that has repeatedly been observed (Mable, Dart, Berardo, & Witham, 2005; Charlesworth, 2006). Moreover, the potential benefits of selfing may be counteracted by inbreeding depression, that is, fitness reduction of selfed progeny in relation to that of outcrossed progeny (Lande & Schemske, 1985). In species with repeated selfing, purging effects may eventually lead to decreased inbreeding depression (Crnokrak & Barrett, 2002); as a consequence, the numerical transmission advantage (3:2) of selfers can drive their increase in populations (Fisher, 1941; Barrett, 2010; Busch & Delph, 2011). However, mixed-mating taxa can have inbreeding depression rates as great as those for outcrossing taxa, indicating that allele purging does not always occur (Winn et al., 2011). In some species with mixed-mating systems, high levels of inbreeding depression hinder the recruitment of selfed progeny, thus maintaining high genetic diversity in the population (Michalski & Durka, 2007). However, in others, reproductive assurance benefits override the inbreeding depression detriment, and plants of selfed origin are maintained in populations despite their low genetic diversity (Michalski & Durka, 2007; Arista et al., 2017). Thus, in conditions of pollen limitation, the reproductive assurance benefits of selfing could be selected (Schoen & Brown, 1991; Cheptou & Massol, 2009) and it can be an important mechanism in maintaining polymorphisms (Narbona et al., 2018) if inbreeding depression is not too high and selfed plants reach reproduction.

To the best of our knowledge, the role of selfing as a factor maintaining flower colour polymorphism has been described exclusively in *Ipomoea purpurea* (Fry & Rausher, 1997; Rausher, Augustine, & VanderKooi, 1993; Subramaniam & Rausher, 2000). In this species, insects stop visiting a floral phenotype when its frequency is very low; these plants then produce seeds through automatic self-pollination, which increases their frequency in the next generation (Subramanian & Rausher, 2000). However, given that more than 40% of species have mixed-mating systems (Goodwillie, Kalisz, & Eckert, 2005; Shivanna, 2015) it is very likely that other polymorphic species show reproductive assurance, this being a neglected mechanism to maintain polymorphisms (Narbona et al., 2018). Nonetheless, consistent differences in mating systems between colour morphs should contribute to reproductive isolation and could initiate a speciation process (Nosil, Harmon, & Seehausen, 2009). Flower colour is considered a trait that promotes speciation by contributing to assortative mating between morphs (Servedio, Van Doorn, Kopp, Frame, & Nosil, 2011). Thus, colour polymorphism can promote the generation of new species; this process ends with the loss of that polymorphism (Hugall & Stuart-Fox, 2012).

Here we study the role of selfing capacity in maintaining flower colour polymorphism in *Lysimachia arvensis*. This species has plants with red or blue flowers that show a geographical pattern of morph distribution; the blue morph is associated with the more stressed and dry zones of the Mediterranean Basin and the red with more temperate areas (Arista et al., 2013). In the Mediterranean Basin, most populations are monomorphic blue or blue-biased. Small solitary bees, the main pollinators of this species, discriminate clearly between flower morphs and show a strong preference for blue-flowered plants in Mediterranean populations (Ortiz et al., 2015); no information about pollinator attendance in non-Mediterranean areas exists. In the Mediterranean, directional selection driven by pollinators gives rise to higher male and female fitness of the blue morph relative to the red morph (Ortiz et al., 2015). Flowers of both morphs show lateral and vertical herkogamy, sequentially (Jiménez-López, Arista, Talavera, Pannell, & Ortiz, 2019), but they differ in the level of each herkogamy type and can self-pollinate autonomously at different times in their lifespan. In the red morph, self-pollen deposition is possible at the end of the flower lifespan (delayed selfing), allowing reproductive assurance if outcross pollination fails (Jiménez-López, unpub. data). Despite the fact that both abiotic and biotic factors negatively select for the red morph in the Mediterranean, it is maintained in a low proportion in these populations. This maintenance suggests the existence of a possible mechanism protecting the red morph. We hypothesize that selfing confers reproductive assurance to the red morph when pollinator attendance is low, thus maintaining it in Mediterranean polymorphic populations. Alternatively, the maintenance of the red morph in the Mediterranean could be driven by gene flow from temperate areas although the small population sizes and the lack of seed dispersal mechanism hardly support this possibility. If selfing is maintaining red-flowered plants in Mediterranean areas, their genetic diversity in Mediterranean populations should be lower than that of blue-flowered plants, and inbreeding depression should be low enough to allow plant recruitment. Thus, we first analysed experimentally the impact of inbreeding depression trough the whole life cycle in two Mediterranean sites where both morphs co-occur. Secondly, we characterized the genetic variation and genetic distance of 20 natural populations occurring in Mediterranean and non-Mediterranean areas taking into account flower colour. Selfing reduces gene flow among individuals (Michalski & Durka, 2007); hence, it could be an isolation mechanism leading to speciation (Martin & Willis, 2007; Brys, Broeck, Mergeay, & Jacquemyn, 2014). If the red morph reproduces predominantly by selfing, this fact could contribute to its reproductive isolation and ecological divergence from the blue morph, and one could expect a genetic divergence of morphs as different evolutionary lineages, or even as incipient species.

## Material and Methods

### STUDY SPECIES

*Lysimachia arvensis* (L.) U. Manns & Anderb. (former *Anagallis arvensis* L.) is a self-compatible annual forb which offers only pollen as a reward for pollinators. It inhabits cultivated fields, wastelands and coastal sands (Ferguson, 1972), and is native to Europe and the Mediterranean Basin; however, it is widely distributed over a large part of the world (Pujadas, 1997). Flowers are visited by small solitary bees that build their nest in the soil near the plants. The fruit is a capsule and the seeds are dispersed by gravity mainly under the mother plants. In general populations are very small. In Mediterranean areas, populations in which the two morphs coexist are frequent, although the blue morph is usually found in a higher proportion (Arista et al., 2013). In contrast, in the Atlantic or temperate areas of Europe, monomorphic red populations are the norm.

### MOLECULAR ANALYSES

AFLP and nuclear microsatellite markers were used to assess how genetic variation is structured among and within populations and floral colour morphs of *L. arvensis*. To this end, 20 natural populations were sampled: 14 from the Mediterranean Basin and six from Non-Mediterranean areas. Of these, 11 were polymorphic and nine monomorphic (four blue and five red; Table 1). Voucher specimens of each population were deposited at the Herbarium of the Seville University (SEV). Six plants of each colour morph were sampled in polymorphic populations and ten in monomorphic populations; leaves of these plants were dried in silica gel and stored until molecular analyses were made. Total genomic DNA was extracted from dry leaf tissue with a plant extraction kit (Invisorb Vegetal DNA Kit HTS 96, Invitek, Berlin, Germany) following the supplier’s instructions. The average DNA concentration was estimated photometrically using a NanoDrop DS-11 Spectrophotometer (DeNovix).

AFLP analyses were performed according to Vos et al. (1995) on 203 individuals, of which 10% were additionally replicated in order to exclude non-reproducible bands. Total genomic DNA was digested with the restriction enzymes EcoRI/MseI. After digestion, adaptors were ligated on both ends of genomic fragments and a two-step selective amplification was performed. We chose six selective primer pairs: EcoRI (Ned)-AAC/MseI-CTG, EcoRI (Vic)-ACG/MseI-CTG, EcoRI (Fam)-ACT/MseI CAG, EcoRI (Ned)-AGC/MseI CTC, EcoRI (Vic)-ACG/MseI-CAT, and EcoRI (Fam)-ACC/MseI-CTC. Resulting PCR products were separated by capillary gel electrophoresis on an automated sequencer (3730 DNA Analyser, PE Applied Biosystems, Foster City, CA, USA) with an internal size standard (GeneScan 500 LIZ, Applied Biosystems) at STABVIDA Lda. (Oeiras, Portugal). AFLP patterns were visualized with GeneMarker 1.9 (SoftGenetics, State College, PA, USA) for manual scoring of fragments after normalisation of the profiles. A fluorescence threshold set at 100 relative fluorescent units was applied to validate the peaks that exceeded the fluorescence intensity of this threshold. Amplified fragments from 100 to 500 base pairs were scored and exported as a presence/absence matrix.

**TABLE 1.**
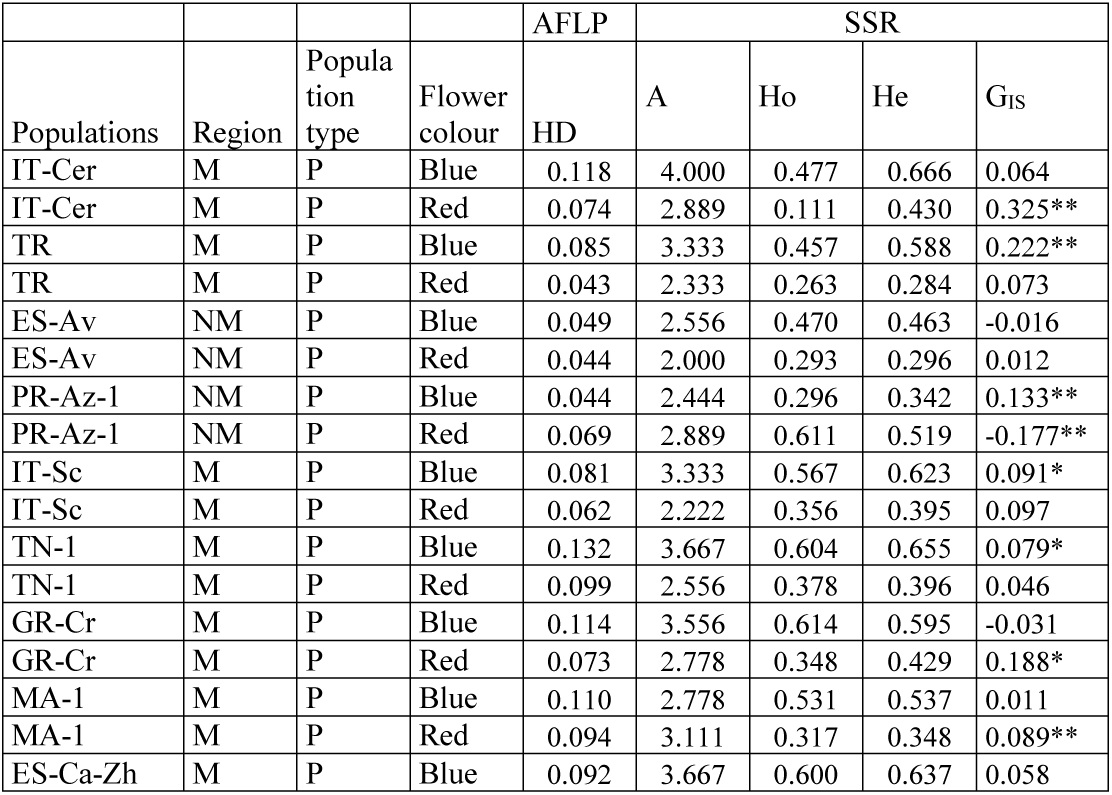
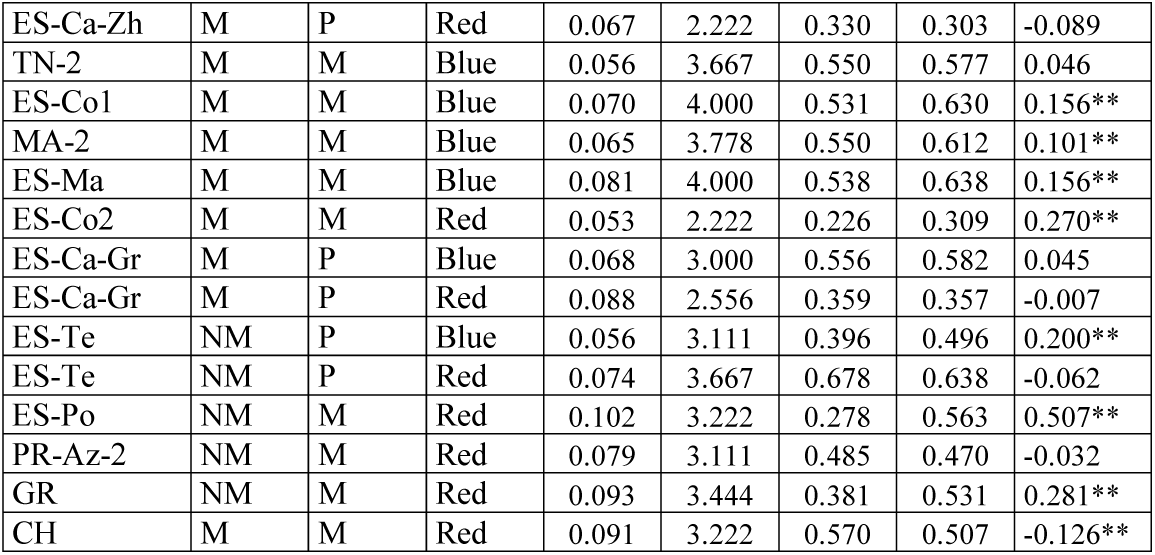
Gene diversity estimates for AFLP and mean of nine SSR microsatellites for each population and colour morph of *Lysimachia arvensis*. Measurements were taken in six red-flowered and six blue-flowered plants in polymorphic populations and in ten plants in monomorphic populations. Region (M, Mediterranean; NM, Non-Mediterranean). Population type (P, polymorphic; M, monomorphic). HD, gene diversity; A, Allele number per locus; Ho, observed heterozygosity; He, expected heterozygosity; G_IS_, inbreeding coefficient (**p<0.01; *p<0.05).

The same 203 individuals were analysed at nine microsallite loci (*Lys*11, *Lys*12, *Lys*16, *Lys*28, *Lys*29, *Lys*30, *Lys*31, *Lys*32 and *Lys*33) previously characterized and available for *Lysimachia arvensis* (Jiménez-López, Talavera, Ortiz, & Arista, 2015). PCR products produced clear amplifications of the expected size on agarose gels. The amplification products were separated by capillary gel electrophoresis on an automated sequencer (3730 DNA Analyser, PE Applied Biosystems, Foster City, CA, USA) with an internal size standard (GeneScan 500 LIZ, Applied Biosystems) at STABVIDA Lda. (Oeiras, Portugal). The scoring was carried out with GeneMarker 1.9 (SoftGenetics, State College, PA, USA) following the same procedure as for AFLP markers.

Gene diversity was estimated separately with data from each molecular marker, for blue vs red morphs and for Mediterranean vs non-Mediterranean populations. From AFLP data, gene diversity was calculated as average gene diversity (HD) with GENEPOP v4.0.10 (Raymond & Rousset, 2011). From SSR microsatellites data, gene diversity was calculated as expected heterozygosity (He) with GENODIVE, version 2.0b25 (Meirmans & Van Tienderen, 2004), and observed heterozygosity (Ho) with ATETRA version 1.2 (Van Puyvelde, Van Geert, & Triest, 2010). Allele number (A), and inbreeding coefficient (G_IS_) were also calculated with GENODIVE 2.0b25, assuming infinite alleles and corrected for unknown allele dosage. Linkage disequilibrium (LD) was calculated with genetics R package (Warnes & Leisch, 2005) after diploidization of each locus, and null allele frequency (No) was estimated by POLYSAT (Clark & Jasieniuk, 2011). To discard the possibility that gene flow maintains the red morph in the Mediterranean, isolation-by-distance between populations of each color morph was investigated by computing the correlation between the matrix of pair-wise population genetic distance obtained by both AFLP and SSR (Φ_PT_) and the matrix of geographical distances, by applying the Mantel test (10000 permutations). Given that *L. arvensis* lacks dispersal mechanism, gene flow is constrained by geographical distance and it would be much more likely to occur between neighbouring populations, following a stepping-stone model.

### INBREEDING DEPRESSION THROUGHOUT LIFE CYCLE

Inbreeding depression (ID) at different stages of the life cycle was studied for both colour morphs under natural field conditions. Given that inbreeding depression depends on the context, we selected two Mediterranean sites where both morphs co-occur: Dos Hermanas and Sevilla. Both sites consist mainly of herbaceous communities on wastelands around orchards. Seeds from plants of both colours were collected in the field, germinated and grown in glasshouses. For each separate colour morph, hand self- and cross-pollinations were carried out to obtain selfed and crossed seeds. The number of seeds per fruit after selfing and outcrossing was recorded in 40-110 fruits of each cross type and colour morph (hereafter seed production of mother plants). A total of 1538 selfed and 1507 outcrossed seeds were sown, 1366 seeds were from blue plants and 1679 from red ones. In the two natural sites, seeds were placed in individual cardboard pots and each potted seed was treated as an independent experimental unit. Sowing was carried out at the beginning of November, and potted plants were harvested at the end of May. During the growth cycle pots were checked every fortnight, and time from sowing to germination (hereafter time to germination), seed germination, seedling survival up to reproductive age, time from germination to flowering (hereafter time to flowering) and seed production after free pollination (hereafter seed production of progeny) were recorded for each plant. Seed production of progeny was estimated as the mean number of seeds in two ripe fruits per plant.

Partial ID coefficients (δ_i_) were calculated for each colour morph and site at each of the following life stages: seed production of mother plants (δ_sm_), total seed viability (δ_tv_), seedling survival (δ_ss_), and seed production of progeny (δ_sp_). To avoid bias in the assessment of inbreeding depression from germination data due to seed dormancy, a subset of selfed and outcrossed seeds was sown in Petri dishes and placed in a germination chamber. Non-germinated seeds (204 self red, 198 cross red, 168 self blue and 168 cross blue) were placed in a 100μl solution of tetrazolium 0.11% to determine their viability (Glenner, 1990). Data regarding germination and viability of non-germinated seeds was then considered together to calculate inbreeding depression at that stage (total seed viability, δ_tv_). Partial ID coefficients were calculated using the expression proposed by Ågren and Schemske (1993):

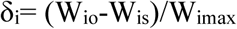

where **δ_i_** is the ID coefficient at life-stage **i**, **W_io_** is fitness after outcrossing, **W_is_** is fitness at this life-stage after selfing, and **W_imax_** is the maximum fitness at this stage (W_io_ or W_is_). If W_io_>W_is_, δ values are positive and inbreeding depression exists at this stage, while in W_is_>W_io_, δ values are negative and outbreeding depression may occur. Cumulative ID coefficients (δ_T_) were also calculated for each colour morph and site. Values of fitness at each life-stage were relativized to the maximum at this stage; cumulative fitness (W_T_) for each cross type (W_To_ and W_Ts_) was then estimated by multiplying relative fitness values at the four life-stages considered (seed production of mother plants, total seed viability, seedling survival and seed production of progeny). In this way, cumulative fitness is presented as a proportion. Then, as for partial coefficients, δ_T_ was calculated by using the expression: δ_T_ = (W_To_-W_Ts_)/W_Tmax_.

### STATISTICAL ANALYSES

To apportion genetic variation as estimated from both molecular markers among and within populations and floral colour morphs, multi-locus analyses of molecular variance (AMOVA) were performed using Arlequin (Schneider, Roessli, & Excoffier, 2000). These analyses hierarchically partitioned molecular variation into within- and among-population components to estimate genetic structure in the following predefined groups: blue vs red plants, Mediterranean vs non-Mediterranean, blue Mediterranean vs red Mediterranean, blue non-Mediterranean vs red non-Mediterranean. Permutation tests were used to determine statistical significance (Excoffier, Smouseand, & Quattro, 1992). For SSR, statistics for the significance (OSx-statistic; Goudet, 1995) across all groups or between pairs of comparison were obtained by 9999 randomizations using GENODIVE 2.0b25.

In exploring the possibility of inbreeding depression, differences in seed production of mother plants and viability of non-germinated seeds were analysed with colour and cross-type (selfing or outcrossing) as fixed factors and taking their interaction into consideration. In analysing differences in time to germination, seed germination, seedling survival, time to flowering and seed production of progeny, the factors site, cross type and colour were considered fixed, and three-way interaction was also considered. GLMs were carried out with different link functions and error distributions, depending on the type of response variable modelled. Binomial distribution of errors and logit link function were used to analyse germination, viability and survival. Binomial negative distribution with log link function was used to analyse time to germination, and normal distribution to analyse time to flowering and seed production. All these analyses were carried out using the GLM module of SPSS (IBM SPSS Statistic 23, 2015, USA) with Type III tests. When GLMs showed significant differences, the means of treatment were compared using t-tests based on standard errors calculated from the specific model.

## Results

### PROPERTIES OF AFLPS AND MICROSATELLITES

The total number of AFLP bands found among populations of *L. arvensis* was 870, of which 82.9% were polymorphic. All individuals had unique AFLP phenotypes and the reproducibility of the AFLP bands was 10%. The nine microsatellite loci were successfully genotyped in the 203 individuals of *L. arvensis*. There were cases of deviation from HWE (*P* < 0.05) after Bonferroni correction across populations and loci; the most significant cases were related to negative or high levels of *G*_IS_, indicating HWE deviation caused by heterozygote excess. Null allele frequency (No) estimated using POLYSAT resulted in the highest frequency of 0.478 (IT-Sc-B/Lys12), with an average frequency of 0.138 over all of the nine markers. Significant LD was not found between any pairwise combinations of loci (*p* < 0.05) after Bonferroni correction.

### GENE DIVERSITY AND POPULATION STRUCTURE

In a total of 203 plants and nine SSR analysed, the total number of alleles was 74 and the frequency of alleles per locus by population and morph ranged from 1.602-2.925. Gene diversity estimates varied between markers; the lower values were found in AFLP (HD) ranging from 0.043-0.118 (mean 0.078 ± SD 0.022) while for multi-locus SSR data, expected heterozygosity (He) ranged from 0.283-0.666 (average 0.497 ± SD 0.027; Table 1). Per locus, He ranged from 0.000-0.862, observed heterozygosity (Ho) from 0.000-0.900 and the inbreeding coefficient (G_IS_) from −0.800 to 1.000. Positive values of G_IS_ were found in the nine loci, indicating heterozygote deficiency (Table 1).

For AFLP, blue plants showed higher gene diversity (0.080 ± 0.006) than red ones (0.065 ± 0.006). For SSR, the blue morph showed significantly higher observed heterozygosity (Ho) values and lower inbreeding coefficient (G_IS_) than the red morph (Ho: 0.562 blue vs 0.415 red, G_IS_: 0.330 blue vs 0.618 red; p<0.05 in all cases). However, the expected heterozygosity was similar between morphs (He: 0.657 blue vs 0.650 red; p>0.05). Taking areas into account, blue plants showed higher gene diversity for AFLP (0.093 ± 0.006) than red ones (0.065 ± 0.006) in Mediterranean areas, but the opposite pattern was observed in non-Mediterranean populations (blue 0.043 ± 0.011; red 0.064 ± 0.008). For SSR, the blue morph showed significantly higher observed and expected heterozygosity and lower inbreeding coefficient than the red morph in Mediterranean areas; in contrast, in non-Mediterranean areas the observed and expected heterozygosity were statistically similar between morphs, but the inbreeding coefficient was statistically higher in the blue morph (Table 2). According to the Mantel tests neither blue-flowered plants nor red-flowered plants showed any association between genetic distance (obtained by AFLP or SSR) and the geographical distance between pairs of populations (for the blue morph, R = 0.144, P = 0.245 by AFLP and R=0.145, p= 0.226 by SSR; for the red morph R =0.101, P = 0.283 by AFLP and R=0.185, p= 0.131 by SSR). Therefore, the hypothesis of isolation by distance was rejected in both color morphs.

**TABLE 2.** Mean values (standard error) of gene parameters for blue and red morphs of *L. arvensis* at nine SSR loci in Mediterranean and non-Mediterranean populations. Within each column, means followed by the same letter are statistically similar. * p<0.05

| Type of population | Colour morph | Observed Heterozygosity (Ho) | Expected Heterozygosity (He) | No. of alleles per locus | Inbreeding Coefficient (Gis) |
| --- | --- | --- | --- | --- | --- |
| Mediterranean | Blue | 0.560 (0.039)a | 0.611 (0.030)a | 6.33 (0.745)a | 0.116* (0.034)a |
|  | Red | 0.327 (0.066)b | 0.381 (0.054)b | 6.00 (0.866)a | 0.204* (0.057)b |
| Non-Mediterranean | Blue | 0.388 (0.073)c | 0.434 (0.045)c | 4.44 (0.503)b | 0.141* (0.043)ab |
|  | Red | 0.486 (0.055)c | 0.499 (0.036)c | 5.78 (0.722)ab | 0.086* (0.040)c |

AMOVA consistently demonstrated a significant population structure separating blue plants from red at both AFLP and SSR markers (Table 3). Mediterranean populations were also differentiated from non-Mediterranean at both markers, although the proportion of explained variance was low. Taking into account flower colour, there was significant differentiation between blue and red Mediterranean plants at both SSR and AFLP markers; 59.59% of variance at SSR and 25.31% at AFLP were due to differences between colours. In non-Mediterranean areas blue and red plants also showed genetic differentiation, the variance attributed to flower colour at SSR and AFLP markers being 52.37% and 36.50%, respectively (Table 3).

**TABLE 3.**
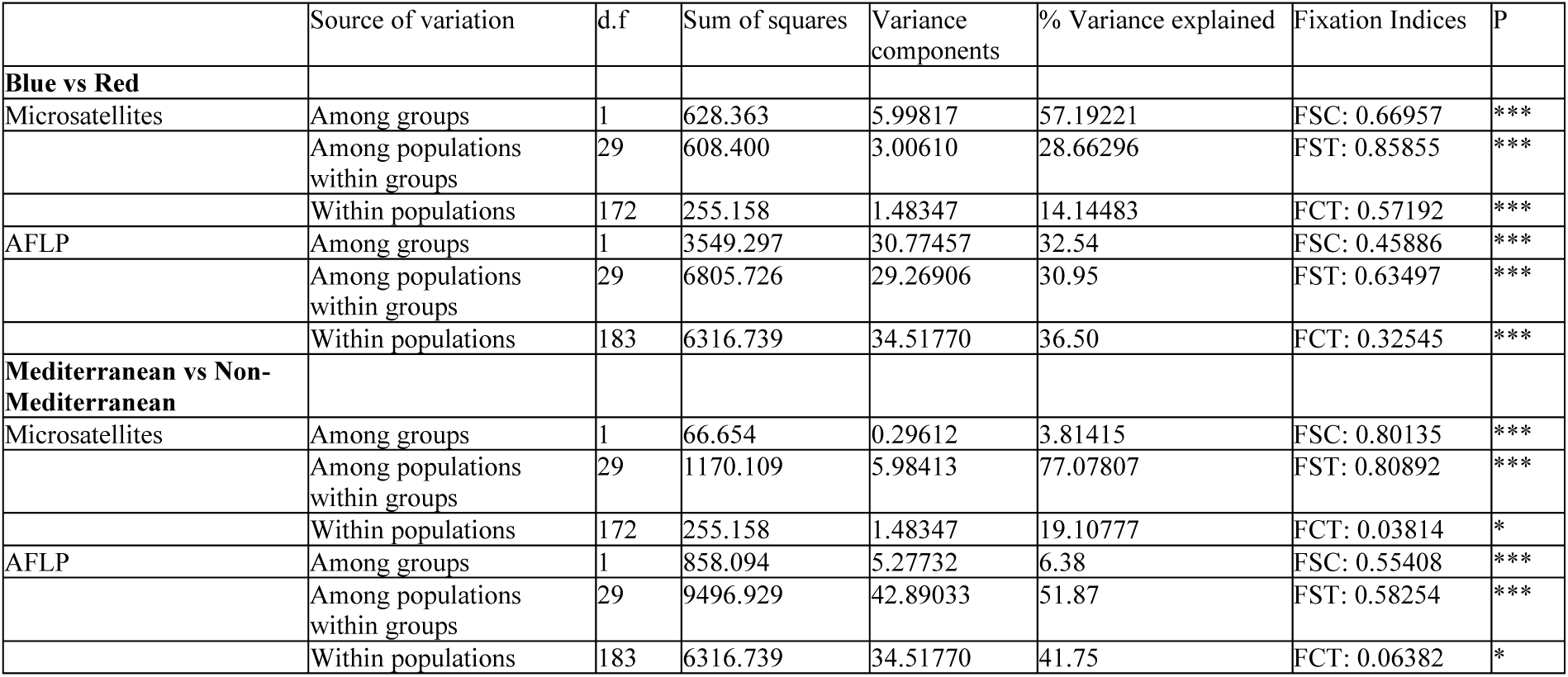
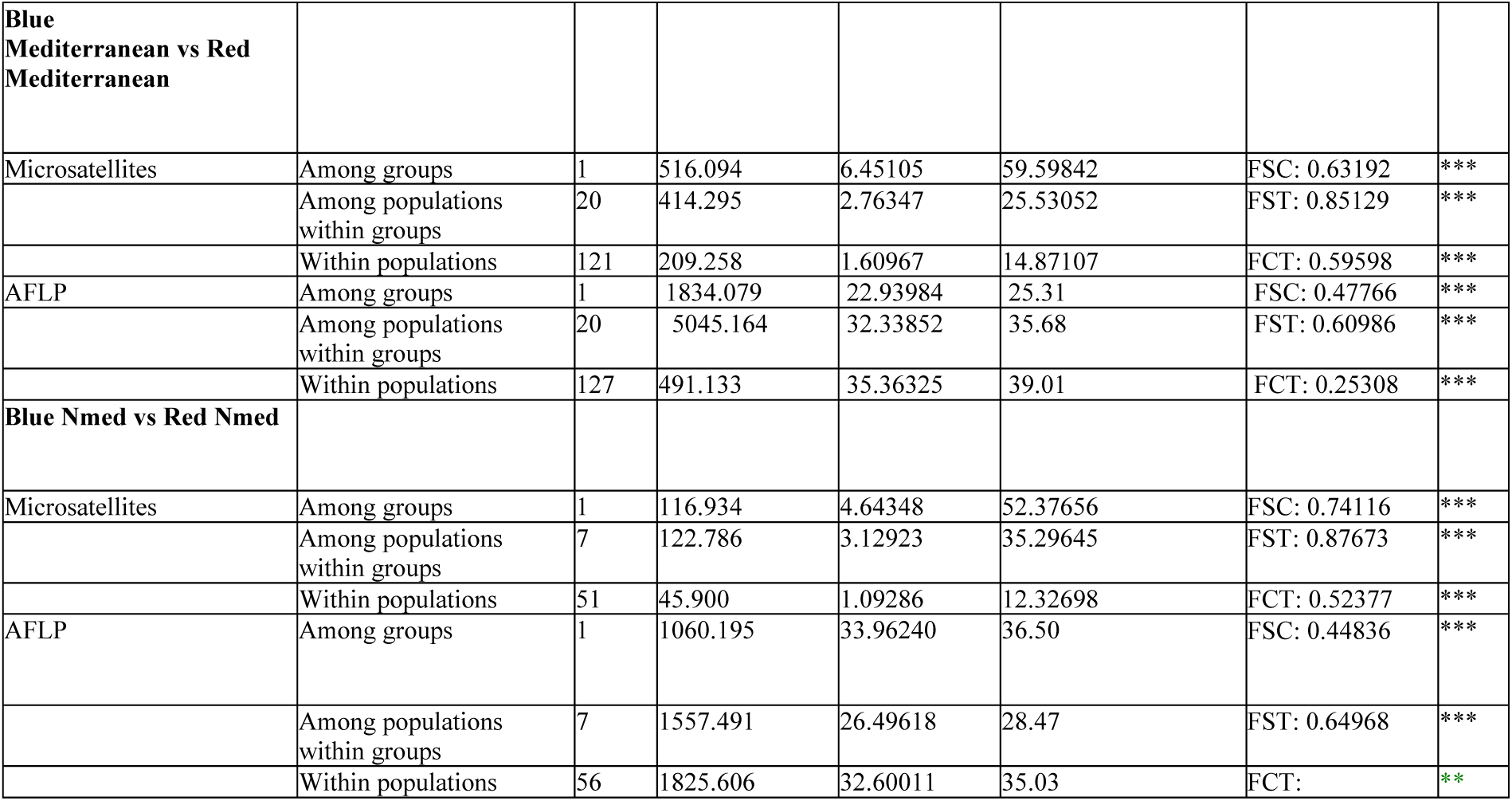
Results of analyses of molecular variance (AMOVA) for AFLP and SSR markers and for different groupings of populations. For microsatellites a total of 9 loci were used. Statistical significance is based on 10100 permutations. * p<0.05, ** p<0.01, ***p<0.001.

### INBREEDING DEPRESSION THROUGHOUT LIFE-CYCLE

The number of seeds per capsule produced after hand-pollination of mother plants differed between treatments (self and cross) and colours (Table 4). In general, seed production was higher in the red morph than in the blue one, and it was also higher after selfing than after outcrossing (Fig. 1). However, the colour-by-treatment interaction was significant (Table 4), as only in the blue morph was seed production significantly higher after selfing than after outcrossing (Fig. 1). Thus, at this first stage of the life cycle, ID coefficient was negative for both colour morphs, although it was very close to zero for the red morph (Table 5).

**TABLE 4.** Summary of GLM results for different effects: site (Sevilla/Dos Hermanas), treatment (selfing/outcrossing) and colour morph (blue/red) and their interactions, on different traits measured in *Lysimachia arvensis*. Significant values in bold.

| Dependent variable | Effects | Wald chi-square | Df | *P* |
| --- | --- | --- | --- | --- |
| Seed production of mother plants | Treatment | 16.211 | 1 | **0.000** |
|  | Colour | 20.335 | 1 | **0.000** |
|  | T x C | 9.232 | 1 | **0.002** |
| Time to germination | Site | 18.097 | 1 | **0.000** |
|  | Treatment | 12.31 | 1 | **0.000** |
|  | Colour | 18.91 | 1 | **0.000** |
|  | P x T x C | 7.85 | 4 | 0.097 |
| Germination | Site | 11.631 | 1 | **0.001** |
|  | Treatment | 16.286 | 1 | **0.000** |
|  | Colour | 26.709 | 1 | **0.000** |
|  | P x T x C | 90.022 | 4 | **0.000** |
| Viability of non-germinated seeds | Treatment | 191.231 | 1 | **0.000** |
|  | Colour | 0.884 | 1 | 0.347 |
|  | T x C | 1.003 | 1 | 0.317 |
| Seedling survival | Site | 1.61 | 1 | 0.204 |
|  | Treatment | 2.06 | 1 | 0.151 |
|  | Colour | 34.58 | 1 | **0.000** |
|  | P x T x C | 24.32 | 4 | **0.000** |
| Time to flowering | Site | 89.42 | 1 | **0.000** |
|  | Treatment | 286.60 | 1 | **0.000** |
|  | Colour | 170.93 | 1 | **0.000** |
|  | P x T x C | 46.88 | 4 | **0.000** |
| Seed production of progeny | Site | 0.70 | 1 | 0.400 |
|  | Treatment | 52.48 | 1 | **0.000** |
|  | Colour | 76.74 | 1 | **0.000** |
|  | P x T x C | 8.16 | 4 | 0.080 |

**TABLE 5.** Estimates of inbreeding depression for the two colour morphs of *Lysimachia arvensis* in two natural sites. Wo and Ws are fitness measures after outcrossing and selfing, respectively, at different life stages (seed production of mother plants, number per fruit; total seed viability, %; seedling survival, %; and seed production of progeny, number per fruit) or total for the life cycle (cumulative, proportion); δi is the inbreeding depression coefficient at each of the stages or its cumulative value.

|  | Seed production of<br>mother plants |  | Total seed viability |  | Seedling survival |  | Seed production<br>of progeny |  | Cumulative |  |
| --- | --- | --- | --- | --- | --- | --- | --- | --- | --- | --- |
|  | Blue | Red | Blue | Red | Blue | Red | Blue | Red | Blue | Red |
| Sevilla |  |  |  |  |  |  |  |  |  |  |
| $W_o$ | 13.49 | 21.37 | 85.12 | 89.12 | 46 | 35 | 23.84 | 21.48 | 0.44 | 0.95 |
| $W_s$ | 20.87 | 22.40 | 32.85 | 57.50 | 67 | 28 | 22.54 | 15.37 | 0.36 | 0.37 |
| $\delta i$ | -0.35 | -0.05 | 0.61 | 0.35 | -0.31 | 0.20 | 0.05 | 0.28 | 0.18 | 0.61 |
| Dos Hermanas |  |  |  |  |  |  |  |  |  |  |
| $W_o$ | 13.49 | 21.37 | 82.00 | 90.14 | 62 | 49 | 25.31 | 19.98 | 0.65 | 0.95 |
| $W_s$ | 20.87 | 22.40 | 62.60 | 54.10 | 44 | 36 | 20.68 | 15.33 | 0.44 | 0.34 |
| $\delta i$ | -0.35 | -0.05 | 0.24 | 0.40 | 0.29 | 0.27 | 0.18 | 0.23 | 0.32 | 0.65 |

**Figure 1.**
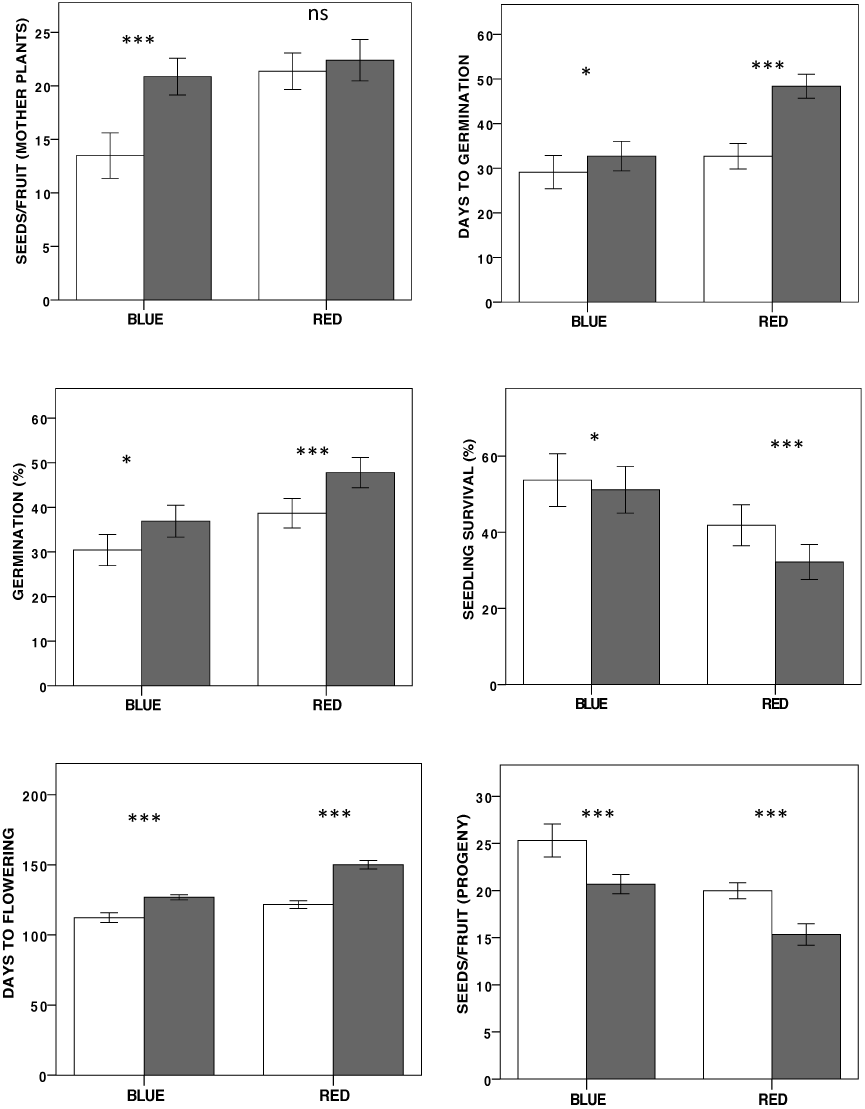
Differences in whole-life fitness components between selfed and outcrossed progeny from the blue and red morphs of *Lysimachia arvensis* in the Mediterranean. Means ±SE are shown. In each graph, asterisks indicate significant differences between selfing and outcrossing values for each colour morph. ***, p<0.001, **, p<0.01, *, p<0.05, ns. Not significant.

In the year studied, two rainy periods occurred and seed germination took place mainly thereafter (Fig. 2). Germination response differed between treatments, colours and sites, but three-way interaction was not significant (Table 4). In general, seeds germinated earlier in the Sevilla site (30 vs. 39 days), and outcrossed seeds germinated earlier in both sites (31 vs. 38 days; Fig. 2). Moreover, seeds from the blue morph germinated more quickly than those from the red morph (mean 29 vs. 39 days respectively).

**Figure 2.**
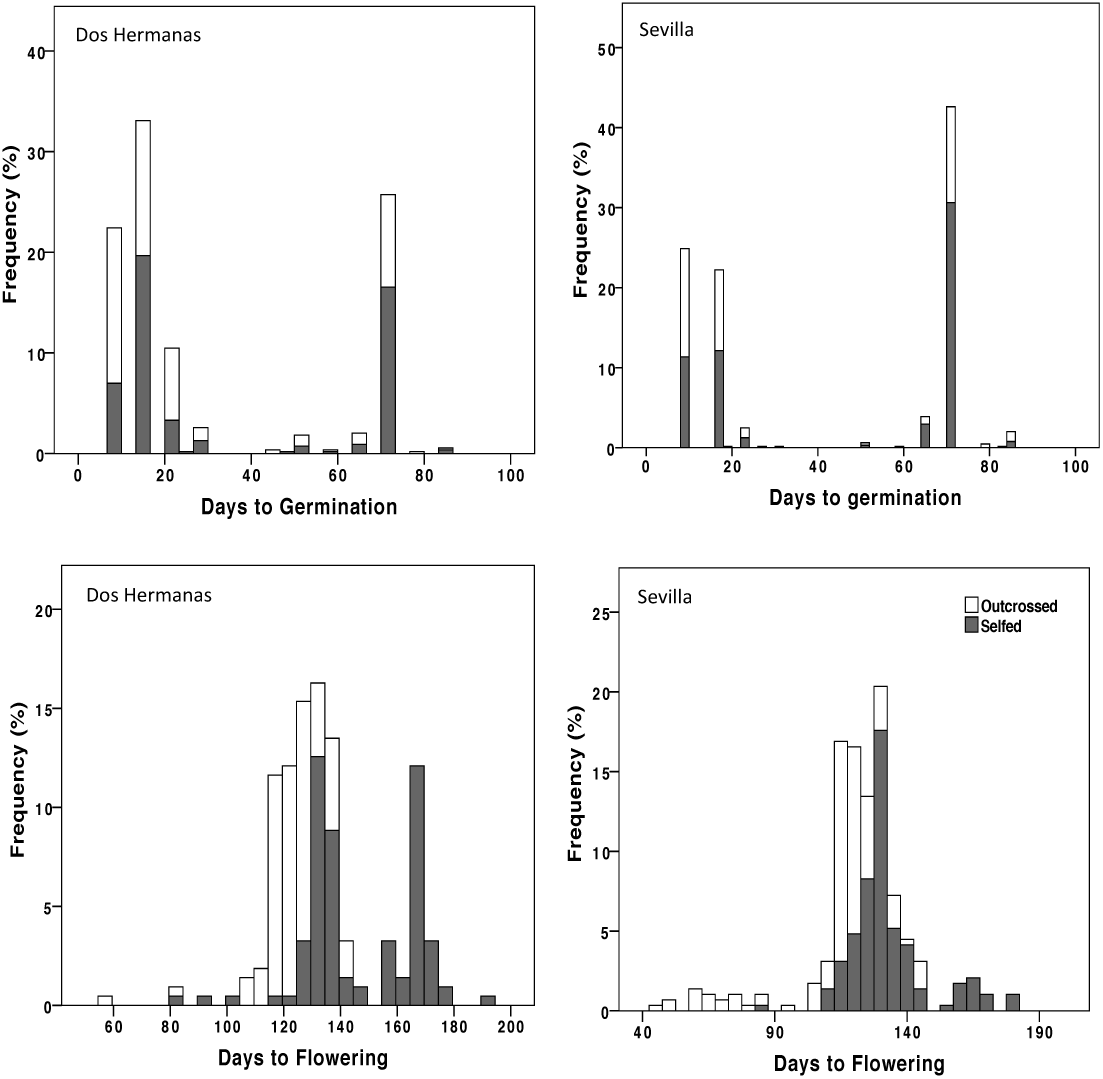
Frequency distributions for two life cycle traits: time to germination and time to flowering of selfed and outcrossed progeny of *Lysimachia arvensis* in two natural sites.

The percentage of seed germination also differed between treatments, colour and sites (Table 4). In general, germination was higher in Dos Hermanas than in Sevilla, in red plants than in blue ones, and in selfed than in outcrossed seeds. However, three-way interaction was significant as the germination patterns were not congruent among sites. Seed viability of non-germinated seeds only differed between treatments (Table 4); most non-germinated outcrossed seeds were dormant but viable, while non-germinated selfed seeds were mainly unviable. Taking into account germination and viability of non-germinated seeds, ID coefficient at this stage was high, although it showed contrasting patterns between morphs and sites (Table 5).

Seedling survival showed differences between colours but not between sites or treatments (Table 4). Three-way interaction was significant here, as a different pattern of survival was shown in each site (Fig. 2). ID coefficient was negative for blue plants in Sevilla but positive for the remaining cases, ranging from 0.20-0.29 (Table 5).

Flowering time showed significant differences between sites, treatments and colour morphs (Table 4), the flowering order being as follows: blue outcrossed plants, blue selfed plants, red outcrossed plants and red selfed plants (Fig. 2). These flowering orders appeared in the two sites studied and as a consequence, there were *L. arvensis* plants in flower for almost five months.

In free pollination, the number of seeds per fruit varied between treatments and colour morphs, but not between sites (Table 4). Outcrossed plants produced a mean of 22.6 seeds while selfed plants produced 18.5 seeds. Blue morphs also produced a higher number of seeds per fruit than red morphs (23.09 vs. 10.04); these differences were found in both sites.

Cumulative inbreeding depression measures were positive for both morphs in both sites, and were higher for the red than for the blue morph. The red morph showed a consistent high ID value of around 0.6 in both sites, whereas in the blue morph it varied from 0.19 in Sevilla to 0.36 in Dos Hermanas (Table 5).

## Discussion

Three main results are derived from this study: 1) a high genetic differentiation between colour morphs that strongly suggests reproductive isolation, 2) a lower genetic diversity of the red morph relative to the blue morph, mainly in Mediterranean areas and not related to an isolation by distance pattern, which suggests differences in breeding system between morphs and 3) a higher inbreeding depression rate for the red morph relative to the blue morph that would render recruitment of selfed progeny difficult in natural populations.

### FITNESS DIFFERENCES OF SELFING AND OUTCROSSING PROGENY

Marked differences in the fitness of the progeny derived from selfing and outcrossing were found in both colour morphs throughout their life cycle. Seed production from selfing originated a higher number of seeds than that from outcrossing in both morphs. This result was unexpected, given that a high impact of selfing was found in the remaining steps in the life cycle; also, seed viability was much higher after outcrossing than after selfing. Given that the flowers of *L. arvensis* are very small and that emasculation is difficult without damaging the flower, floral manipulation should be considered as a possible cause of decreased seed production in hand-outcrossed flowers. Seed germination was also higher in selfed seeds, but this was merely a consequence of differences in seed dormancy between selfed and outcrossed seeds. All outcrossed seeds that did not germinate were viable, while ungerminated selfed seeds were dead. Given differences in germination between morphs, this means that seed-banks of *L. arvensis* in the wild should consist of outcrossed seeds from which individuals could be incorporated into the populations every year.

The progeny derived from selfing also showed marked differences in phenology in relation to that derived from outcrossing in both morphs. This was an unexpected finding, because differences in phenology according to breeding system are not usually reported in the literature. Non-dormant seeds of *L. arvensis* germinated just after a rainy period giving rise to pulses in which the germination order was: first outcrossed blue, then selfed blue and outcrossed red, and selfed red last. In annuals, the time of germination is the first major developmental transition influencing all posterior life cycle traits (Finch-Savage & Leubner-Metzger, 2006; Manzano-Piedras, Marcer, Alonso-Blanco, & Picó, 2014). Arid environments such as those in the Mediterranean are characterized by limited and variable rainfall that supplies resources in pulses (Chesson et al., 2004). In these environments, quick germination just after rain permits seedlings to develop deep roots to tolerate water scarcity, thereby increasing survival probability (Schenk & Jackson, 2002). As blue seeds germinated earlier than red ones, differences in survival found between colours could be a result of germination differences. Survival differences between morphs in dry environments were found experimentally in a previous work (Arista et al., 2013); thus this study confirms previous findings. In the red morph, differences in the time of germination between selfed and outcrossed seeds were much more marked than in the blue morph; consequently, differences in survival between selfed and outcrossed seedlings were high.

Flowering phenology was also markedly affected by progeny origin, with blue plants flowering earlier than red ones, and with outcrossed plants flowering earlier than selfed ones. This implies a variation in mating phenology between colour morphs, as was already reported in the greenhouse (Arista el al., 2013) although overlap also occurs. Differences in reproductive timing often have strong fitness consequences, as they promote assortative mating (e.g. Fox & Kelly, 1993; Fox, 2003). Differences in flowering phenology between morphs could limit pollen flow between them in polymorphic populations, as assortative mating within colours is frequent. Even if colour morphs show a temporal overlap in flowering phenology, assortative mating can be much stronger than expected as the chance of mating is reduced (Fox, 2003). Thus, a difference in flowering phenology acts as a prezygotic barrier to gene flow (Martin & Willis, 2007; Botes, Johnson, & Cowling, 2008) and given that prezygotic barriers generally make a greater contribution to reproductive isolation than postzygotic barriers (Lowry, Modliszewski, Wright, Wu, & Willis, 2008; Widmer, Lexer, & Cozzolino, 2009), mating phenology could contribute efficiently to morph isolation in *L. arvensis*. On the other hand, flowering phenology is constrained by both abiotic and biotic factors (Cruz-Neto, Machado, Duarte, & Lopes, 2011), and can strongly influence plant reproductive success (Ollerton & Lack, 1998). In annual plants, an early flowering when water is available permits an extended flowering period, and in seasonal climates such as the Mediterranean, this is advantageous as it assures plant reproduction (Rodríguez-Pérez & Traveset, 2016). In contrast, a late flowering increases the risk of drought and limits vegetative growth and fruit production (Giménez-Benavides, Escudero, & Iriondo, 2007). In fact, we have found that seed production showed the same pattern as flowering phenology, with higher production in plants flowering earlier (outcrossed blue) and lower in plants flowering later (selfed red). Thus, in the Mediterranean sites studied, the very late flowering of selfed red plants is markedly disadvantageous, strongly limiting seed production. However, given that the plants studied grew in natural conditions, differences in seed production between morphs could also be a consequence of differential pollinator attendance to morphs, as reported in some natural Mediterranean populations of *L. arvensis* (Ortiz et al., 2015).

### GENETIC VARIATION BETWEEN COLOUR MORPHS

The analyses of genetic variation in *L. arvensis* at both neutral DNA markers showed a strong partitioning of molecular variation between the red and blue morphs in both Mediterranean and non-Mediterranean areas. Differences between colour morphs explained most variations in both AFLP and SSR markers. Genetic differentiation between colour morphs in the populations studied strongly indicates that gene flow between them is restricted, morphs being to some extent reproductively isolated. This result is supported by two facts: first, differences in flowering phenology found here and in a previous study (Arista et al., 2013) that hinder pollen flow between morphs, and second, differences in pollinator attendance in polymorphic populations where pollinators prefer blue flowered plants and show floral constancy (Ortiz et al., 2015; Jiménez-López et al., unpub. results). In addition, the subtle but consistent differences in herkogamy traits found between morphs (Jiménez-López et al., unpub. data) could also be a consequence of evolutionary divergence owing to morph isolation.

The red morph had a consistently lower genetic diversity, lower observed heterozygosity and lower number of alleles than the blue morph. Plant mating systems have significant effects on genetic diversity (reviewed by Charlesworth & Wright, 2001), with selfers showing much lower diversity than outcrossers. These differences between selfers and outcrossers are expected to be even more pronounced when both kinds of plants co-occur within populations (Glémin et al., 2006). Thus, the set of differences in gene diversity found between morphs suggests a higher selfing rate in the red morph. The strongest differences in gene diversity were found in Mediterranean areas, where moreover the blue morph showed a higher diversity and a lower inbreeding coefficient. In contrast, in non-Mediterranean areas results for the two markers were not congruent; red plants showed higher genetic diversity than blue ones when AFLP was used, but both morphs showed similar diversity with SSR microsatellites. In non-Mediterranean areas, the inbreeding coefficient showed the opposite pattern to that found in Mediterranean areas, since it was higher for the blue morph, although similar to that in the Mediterranean. Differences in pollinator attendance in Mediterranean and non-Mediterranean areas could explain these differences. In a previous study, pollinators showed a marked preference for the blue morph in blue-biased populations and a similar preference for both morphs in balanced populations (Ortiz et al., 2015).

The authors explained that result as a consequence of positive frequency-dependent pollinator attendance, a very common situation in natural populations (Cresswell & Galen, 1991; Smithson & MacNair, 1996). Given that the blue morph is more frequent in Mediterranean areas while the red morph is much more frequent in non-Mediterranean areas (Arista et al., 2013), differential attendance of pollinators could explain differences in inbreeding coefficient between morphs. This would imply that mating system is context-dependent for the red morph, being mainly selfing in the Mediterranean but outcrossing in non-Mediterranean areas. In contrast, the similar inbreeding coefficient of the blue morph in both areas suggests a similar breeding system in both Mediterranean and non-Mediterranean areas, although parameters of genetic diversity were higher in the Mediterranean.

The higher inbreeding coefficient and the low genetic diversity of red plants in the Mediterranean suggest that some selfed progeny is recruited in populations, despite the high values of inbreeding depression throughout the life cycle found in the field. Differences between selfed and outcrossed progeny were found in both morphs, although they varied in intensity. The ranges of inbreeding depression found in *L. arvensis* are in accordance with those of plants with mixed reproductive systems, ranging from 0.2 to 0.8 (Winn et al., 2011). Interestingly, ID in the blue morph was close to that of selfing species (near of 0.2), while red morph ID was close to that of outcrossing (about 0.6). According to Winn et al. (2011) if purging is occurring, ID of mixed-mating species should be closer to that of selfing species as occurs in the blue morph; this suggests an evolutionary trend towards selfing. In contrast, the high ID of the red morph would suggest the existence of a mechanism to avoid purging, limiting evolution towards selfing and maintaining stability of the mixed-mating system in the red morph (Winn et al., 2011).

The high ID rates in the red morph in the two sites studied (about 0.6) mean that only 40% of selfed progeny could be recruited in populations, while outcross progeny recruits 100%. The impact of ID on the fitness of the red morph was markedly high, but selfing could be the sole way to ensure reproduction under pollen limitation.

Importantly, annual plants must produce seeds before dying, and even low quality offspring make some contribution to fitness, defined as the representation of an individual’s genes in progeny generation (Charlesworth, 2006). Moreover, the delayed selfing mechanism of the red *L. arvensis* avoids an extra cost of pollen and seed discounting; consequently, any surviving selfed progeny would provide reproductive assurance (Kalisz, Vogler, & Hanley, 2004) by maintaining the red morph. Given that the red morph is negatively selected by both biotic and abiotic factors in the Mediterranean Basin (Arista et al., 2013; Ortiz et al., 2015), the survival of any selfed progeny can help its maintenance in populations. The high ID limits the numerical advantage of selfing (3:2; revised in Jain 1976) and impedes an increase in the frequency of the red morph in Mediterranean polymorphic populations.

In short, we have found marked differences in the progeny coming from selfing and outcrossing in both morphs, but also between morphs. The earlier germination and flowering of outcrossed relative to selfed progeny, and that of the blue morph relative to the red morph, can represent an important recruitment limitation for red-flowered plants in Mediterranean areas. The low genetic diversity of the red morph is in accordance with a reproductive system based on predominant selfing. Despite this, the red morph showed higher inbreeding depression rates relative to those of the blue one, suggesting a limited capacity for recruitment of the selfed progeny of the red morph. The strong and consistent genetic differentiation between colour morphs in the populations studied indicates a history of gene flow limitation between them. Phenological and genetic differences between morphs found in this study add up to subtle morphological differences in herkogamy traits (Jiménez-López et al., unpub. data), strongly suggesting that the two colour morphs of *Lysimachia arvensis* are, in fact, different lineages.

## Acknowledgements

This work was supported by the European Regional Development Fund (ERDF) and grants from the Spanish MINECO (CGL2012-33270; CGL2015-63827) to M.A. and P.L.O. and to F.J. J-L. (BES-2013-062859). The authors thank Servicios Generales de Herbario e Invernadero de la Universidad de Sevilla, and the DNA Bank Herb SEV. Furthermore, authors thanks Parque del Alamillo de Sevilla for experimental site supplied.

